# Quantitative comparison between aMRI and DENSE for the assessment of brain tissue motion

**DOI:** 10.1101/2023.01.24.525340

**Authors:** Ayodeji L. Adams, Itamar Terem, Allen Champagne, Samantha J. Holdsworth, Jaco J. M. Zwanenburg

## Abstract

**Purpose:** Amplified MRI (aMRI) holds potential for assessing brain tissue motion and strain, using images acquired from readily-available sequences. However, image registration is necessary to extract displacements from the motion-amplified images, which may limit its accuracy. We aimed to separately assess the errors from imperfections in the aMRI amplification, and errors from the registration algorithm, using a semi-synthetic approach.

**Methods:** Ground truth brain tissue motion was derived from smoothed Displacement Encoding with Stimulated Echoes (DENSE) measurements acquired at 7T (8 subjects). Those were then applied to a still sagittal anatomical balanced-SSFP image to obtain a DENSE-animated MRI series to which aMRI (amplification factor 10) was applied. DENSE-amplified MRI series served as a reference (Damp-MRI; amplification factor, 10). Amplified displacements were extracted from aMRI and Damp-MRI using a common registration algorithm. Linear regression was used to estimate the amplification and r^2^ agreement of the amplified displacements relative to the ground truth.

**Results:** The estimated amplification was consistently lower for aMRI-derived displacements (range: [4.9±0.3 5.7±0.3]) than for Damp-MRI measurements (range: [6.7±0.5 7.7±0.5]). Nevertheless, the spatial, temporal and average characteristics of brain tissue motion derived from aMRI were comparable to the ground truth for Anterior-Posterior and Feet-Head displacements: (group averaged r^2^≥0.84), as were the Damp-MRI derived displacements (r^2^≥0.88). When aMRI was applied to in-vivo cine-bSSFP images and compared to the ground truth, the results were less favorable, highlighting the need for artefact-free images.

**Conclusion:** These results strengthen the potential of aMRI as a tool for semi-quantitative assessment of brain tissue motion in disease.

## 1 Introduction

Brain tissue exhibits cardiac-induced pulsatile displacement which may be altered after injury or disease ^1–4^, providing a potential mechanism to better understand the pathological mechanisms underlying these diseases. Additionally, brain tissue strain is derivable from tissue displacement fields^5–7^, which may provide insights into diseases that affect the (visco)elastic properties of brain tissue, or the cerebral small vessels which act as a conduit for cardiac-induced brain tissue strain^8^. Displacement Encoding with Stimulated Echoes (DENSE)^5,9,10^ and amplified MRI (aMRI)^11,12^ have the sensitivity to discern the subtle displacements which are associated with brain tissue motion. DENSE encodes displacements in the phase of the MR signal, thereby providing access to measurements of sub-voxel tissue motion. DENSE may therefore be considered as a method that provides quantitative ground truth displacements of brain tissue motion, although it is currently limited with respect to its clinical availability. aMRI, which enables the visualization of brain tissue motion, can be used to estimate the displacement fields through a post-processing amplification algorithm applied on conventional cine-images, which yields a great potential for assessing pulsatile brain tissue motion on clinically available scanners with readily available sequences.

Despite recent advances in amplified-based motion imaging^11,12^, the aMRI algorithm yields only the deformed images, and not the displacement fields. Thus, image registration is necessary to quantitatively assess displacements originating from the amplified motion images generated by aMRI, which potentially limits its accuracy for quantitative estimations of brain tissue motion. Additionally, the accuracy of the registration algorithm may be further dependent on tissue contrast and image noise and artefacts, possibly adding to the limitations of aMRI. For those reasons, a direct comparison of the displacements derived from DENSE and aMRI is challenging since it is inherently biased by the confounding errors introduced during image registration. Moreover, accurate quantification of tissue strain within an imaging plane critically depends on good estimation of the amplification factor applied to the two orthogonal components of the 2D displacement field extracted from the registration. This introduces an additional variable that affects the strain calculation, which in itself is already inherently noise-sensitive. Altogether, this provides grounds for the independent assessment of errors introduced by the registration process and the aMRI post-processing, as an avenue to advance the use of amplification algorithms in clinical settings, while having a thorough understanding of its limitations.

In this study, we aimed to separately assess the errors induced by image registration limitations, and by potential imperfections in the aMRI amplification algorithm for quantitative assessment of brain tissue motion. This was done by using a semi-synthetic approach, where a common registration algorithm was used to extract displacements from both aMRI and DENSE-amplified images of brain tissue, which were then compared against a known ground truth. We further utilized these displacement measurements to calculate brain tissue strain parameters, as an additional sub-analysis for comparing the results from aMRI.

## 2 Methods

The assessment of potential errors induced by the aMRI algorithm was performed on both a semi-synthetic dataset and an in-vivo dataset. An overview of this assessment is as follows. First, a description of the acquired images, and post-processing of these measurements is provided since these were used to create the data necessary to perform the semi-synthetic analysis. Secondly, the generation of the semi-synthetic dataset is described in depth. Finally, the details of the analysis performed on all the datasets is supplied. Figure 1 contains an overview of the processing steps.

**Figure 1.**
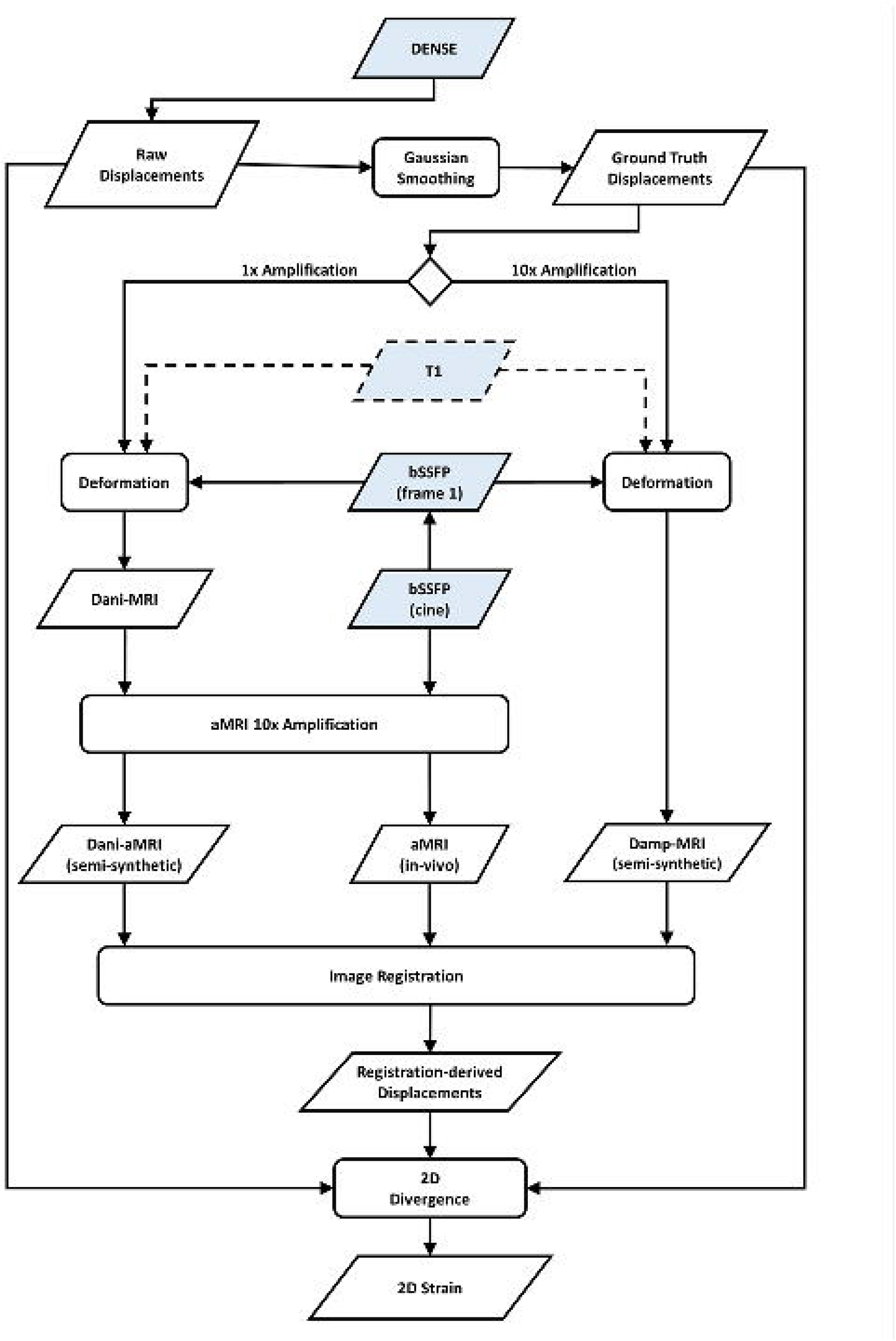
Schematic representation of the processing steps and experiments performed in this study. aMRI = Amplified-MRI, Dani-MRI = DENSE animated MRI, Dani-aMRI = DENSE animated aMRI, Damp-MRI = DENSE amplified MRI. The DENSE, T1-weighted and bSSFP (shaded in blue) were acquired MR images whilst the aMRI, Dani-MRI, Dani-aMRI and Damp-MRI were created using post-processing algorithms. Comparison of Dani-aMRI (registration-derived) displacements and Damp-MRI (registration-derived) displacements to the ground truth displacements allowed for the separate investigation of the errors induced by the registration and aMRI algorithm.

### 2.1 Acquired measurements

A 7T MR scanner (Achieva, Philips Healthcare, Best, The Netherlands) with a 32 channel head coil (Nova Medical, Wilmington, USA) was used to acquire 4D DENSE images with Feet-Head (FH), Anterior-Posterior (AP) and Right-Left (RL) motion encoding directions. Images were acquired as previously described^6^ in 8 healthy subjects of European descent (5 males, mean age: 27±6 years) to extract whole brain tissue displacements covering the entire cardiac cycle. In brief, the following parameters were used: FOV=250×250×190 mm^3^, resolution=1.95×1.95×2.2 mm^3^, motion sensitivity=0.7 mm/2pi (FH) and 0.35 mm/2pi (AP and RL), temporal resolution=51 ms. The scan duration was 3×2.4 min for a heart rate of 60 bpm, for the three orthogonal motion encoding directions (FH, AP and RL).

2D mid-sagittal (10-20 mm left/right from the interhemispheric fissure) balanced-SSFP (bSSFP) cine-images of all subjects were acquired in the same scan session as the DENSE images, using the following imaging parameters: FOV=240×240 mm^2^, resolution=0.83×0.83×3 mm^3^, TR=4.1 ms, TE=2 ms, EPI factor=5 (to achieve a high temporal resolution which is desirable for the aMRI algorithm), number of averages=4, flip angle=20°, cardiac synchronization device=pulse oximeter. The total number of reconstructed cardiac frames was 80, for a total acquisition time of approximately 4 minutes. These bSSFP images served as the in-vivo dataset.

A high-resolution T1-weighted image (FOV=300×248×190 mm^3^, resolution=0.93×0.93×1 mm^3^, TR=4 ms, TE=2 ms, inversion delay=1235 ms, flip angle=5°) was also acquired for all subjects for registration of the processed images in anatomical space.

The governing Ethical Review Board of our institution approved the use of human subjects in this study. Subjects were instructed to lie still in the scanner, and further had their head motion restricted through the use of padding in the transmit/receive head coil.

### 2.2 Post-processing

#### 2.2.1 DENSE

For each subject, the AP and FH DENSE images were first rigidly registered together to correct inter-scan subject motion (with 6 degrees of freedom), and then non-rigidly registered to the T1-weighted image (T1w), resulting in interpolated high-resolution (1 mm isotropic) DENSE images. Registration of the DENSE images with each other, and with the T1w, was performed using the open-source software Elastix^13^. DENSE measurements were temporally adjusted to reflect displacements with respect to end-diastole as reference (vectorcardiogram triggering), instead of at peak-systole (pulse-oximeter triggering). To achieve this (as fully described in Adams et al.^6^), the data were circularly shifted on the time dimension by approximately 250 ms (rounded to an integer number of frames). Subsequent offsets in the displacements were corrected such that displacements at the beginning of the cardiac cycle were zero.

2D ground truth displacements (GTD) were created by spatially smoothing the AP and FH DENSE displacements associated with the same spatial location as the acquired bSSFP images. Smoothing was performed with a Gaussian filter (kernel=21×21 pixels, sigma=3.5). Noise reduction by Gaussian smoothing was found to be a viable method for improving precision, while reducing bias in DENSE measurements^14^. Displacement values outside the brain tissue mask were set to zero before smoothing. The percentage error of the smoothed displacements was determined and found to be small relative to the peak absolute displacement (Figure 2).

**Figure 2.**
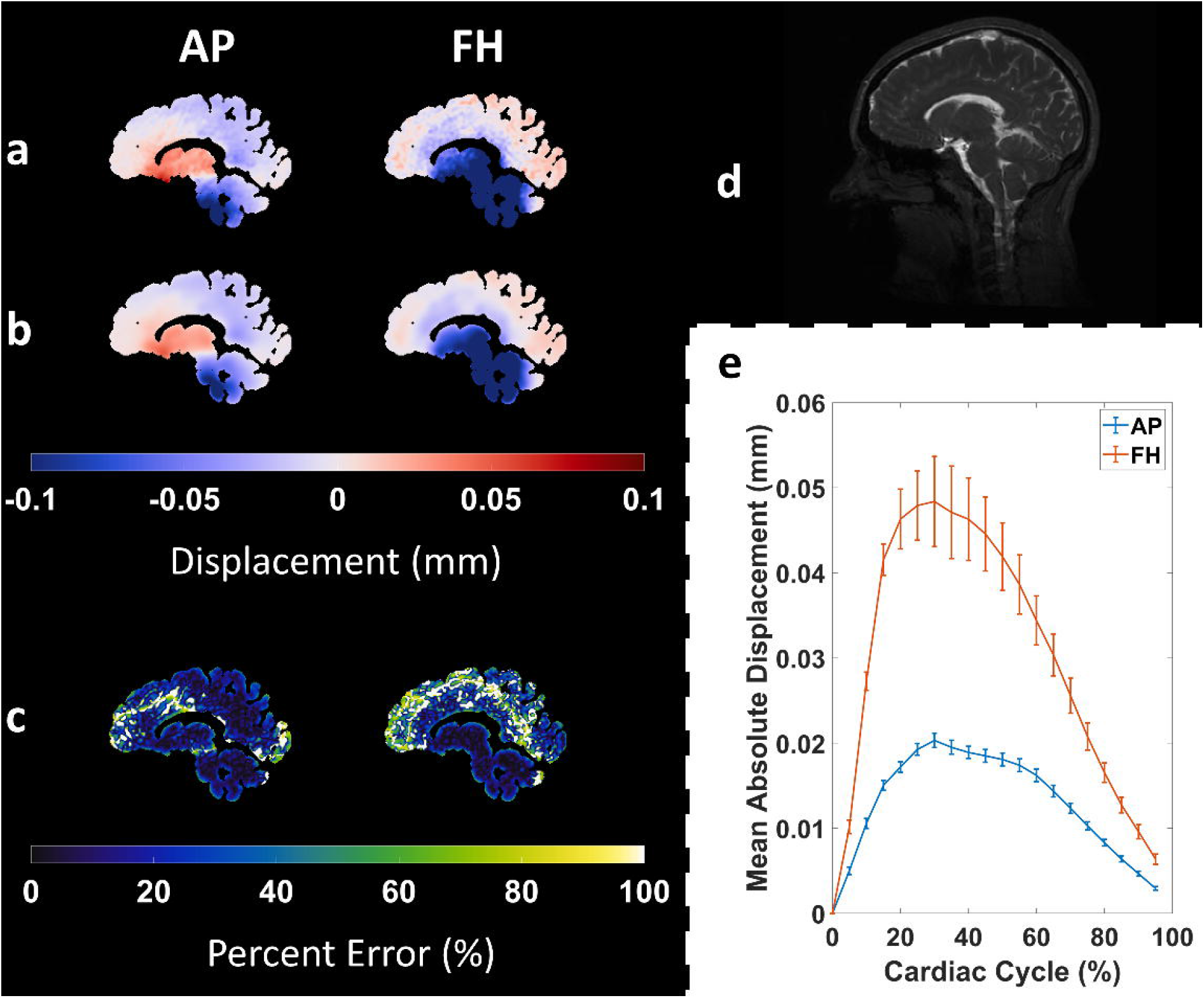
Example AP and FH raw displacements (a) and smoothed ground truth displacements (b) from one subject. Percentage error maps between the raw and smoothed displacements are shown in (c). All maps are shown at the moment of peak displacement for the subject. (d) Reference bSSFP anatomical image. The mean (within the brain tissue) group mean absolute AP and FH ‘raw’ displacements over the cardiac cycle is shown in (e). Error bars represent the group mean absolute difference between the raw and smoothed displacements.

#### 2.2.2 Cine-bSSFP

To facilitate creation of the semi-synthetic data, it was desirable to have the bSSFP images in the same image space as the interpolated high-resolution DENSE. Therefore, 2D mid-sagittal slices were extracted from the high-resolution 3D T1w, at the same spatial locations as the bSSFP images. The bSSFP images were then rigidly registered and transformed to the 2D mid-sagittal T1w space. Registration was performed on the first frame of the bSSFP cine images to obtain the rigid transformation parameters, which were then used to transform all subsequent frames to the T1w anatomical space.

High-frequency artefacts were observed in the bSSFP images (Figure 3). As the aMRI amplification and image registration algorithms are sensitive to intensity fluctuations, the high-frequency components were suppressed using a minimum order, infinite impulse response low-pass filter with a cut-off frequency of 3Hz.

**Figure 3.**
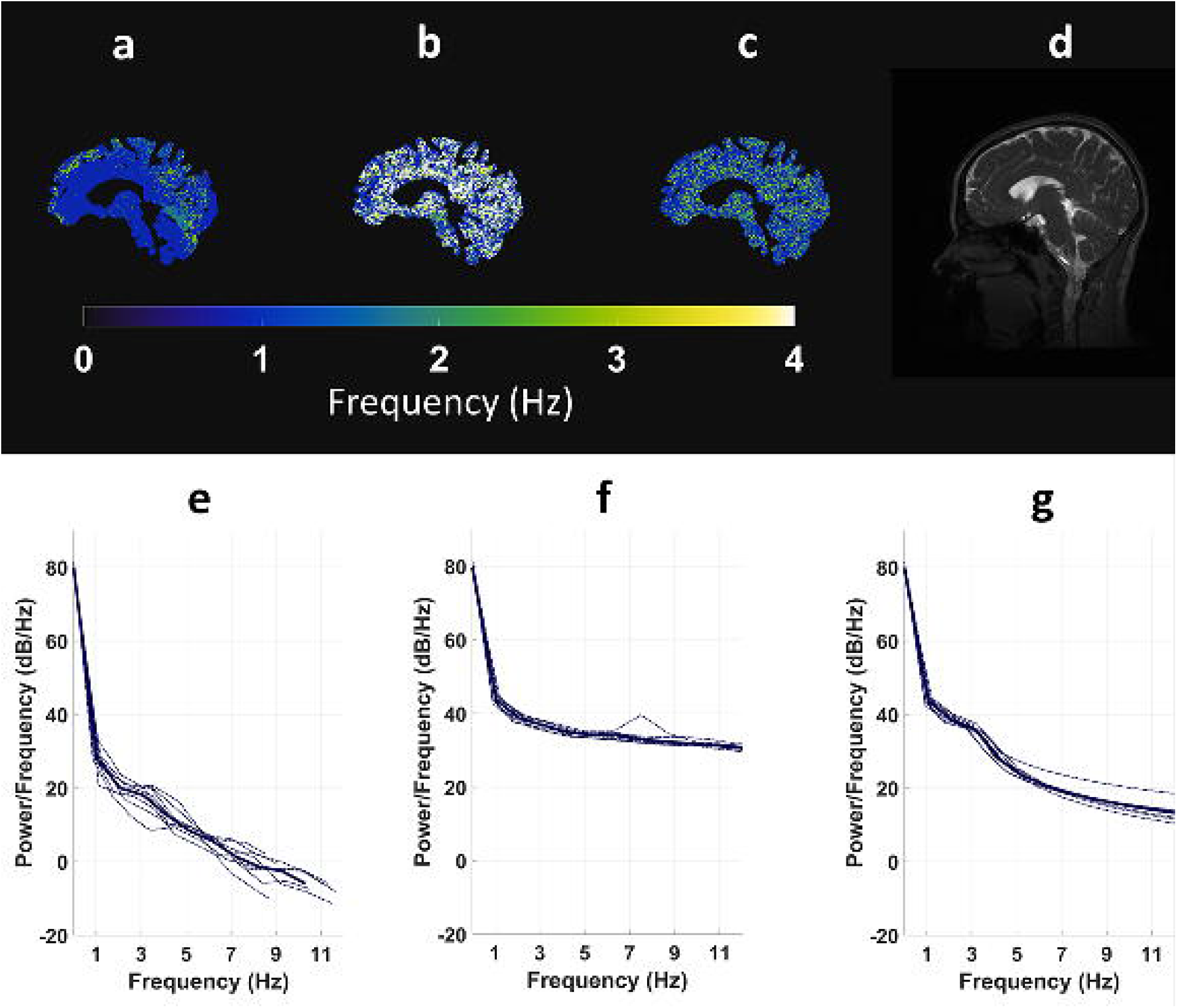
Example max power frequency maps for Dani-MRI (a), the unfiltered in-vivo-aMRI (b) and the low pass filtered in-vivo-aMRI (c). Reference magnitude image is shown in (d). The mean power (with the brain tissue) at different frequencies for the Dani-MRI, the unfiltered in-vivo-aMRI and low-pass filtered in-vivo-aMRI in all subjects is shown in e,f and g, respectively. The bold line signifies the group mean value.

#### 2.2.3 Tissue masks

Brain tissue masks were generated by performing automatic tissue segmentation using the Computational Anatomy Toolbox (Jena University Hospital, Departments of Psychiatry and Neurology) in SPM12 (Wellcome Trust Centre for Neuroimaging, University College London). Voxels containing only CSF, or partial-volumed with CSF (as determined by the segmentation software) were removed from the brain tissue masks. Regions containing DENSE artefacts were further excluded from the brain tissue masks using an SNR threshold as previously described^6^.

### 2.3 Semi-synthetic dataset

Synthetic data were necessary to untangle the combined errors induced by the aMRI and registration algorithms. To create a test environment with realistic brain tissue motion, the DENSE measurements were used to create a synthetic dataset (hereafter referred to as semi-synthetic data). This was accomplished as follows: the displacement fields obtained through DENSE were used to deform (‘animate’) a reference image, thereby providing an animation of the tissue motion over the cardiac cycle. The first frame of the bSSFP acquisition was selected as the reference image, which was deformed using the GTD. The resulting dataset is termed as DENSE-animated images (Dani-MRI). DENSE-amplified images (Damp-MRI) were similarly created by linearly scaling the GTD by a factor of 10 and then using the scaled GTD to deform the bSSFP reference image.

### 2.4 Analysis

The following analysis steps were performed on the semi-synthetic and in-vivo datasets.

#### 2.4.1 Phase-based amplification

The phase-based aMRI algorithm^12^ was used to amplify the motion in the Dani-MRI and the in-vivo bSSFP cine-images. For clarity, we term the amplification of Dani-MRI and in-vivo bSSFP images with aMRI as Dani-aMRI and in-vivo-aMRI, respectively.

aMRI takes as an input a time series of MR images, and outputs images with magnified motion. The main idea behind aMRI is that the motion field is not explicitly estimated, but rather magnified by amplifying temporal intensity changes at a fixed position. Magnification is achieved by decomposing the series of MR images into scale and orientation components using a linear complex-valued steerable pyramid. Temporal variations in the phase of the coefficients of the complex-valued steerable pyramid correspond to motion and can be temporally processed and amplified to reveal imperceptible motion, or attenuated to remove distracting changes.

The original aMRI algorithm^12^ enables one to selectively amplify different temporal frequencies, with different amplification factors depending on the physiological information of interest. In addition, an optional amplitude-weighted Gaussian spatial smoothing can be performed on the filtered phases, in order to increase the phase SNR and support a larger amplification factor. In this study, we wanted to quantify and compare the overall motion, while avoiding noise from higher temporal frequencies which existed in the in-vivo bSSFP data. As such, in addition to the low-pass filtering applied on the cine-bSSFP images, we also chose to selectively amplify motion within the range of 0-3 Hz. The use of a cardiac-gated acquisition ensured that most energy was concentrated within this range^12^. Additionally, in order to avoid the loss of motion information, and also perform a better comparison between the Dani-aMRI and Damp-MRI images, we did not apply the optional amplitude-weighted Gaussian filter on the filtered phases.

Using the displacement function *δ*(*t*) and spatial wavelength *λ*, the amplification parameter a was bounded in the original phase-based study^15^ as 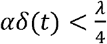 for the regular octave complex steerable pyramid, and 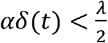 for the sub-octave complex steerable pyramid. In order to support a larger amplification factor, we decided to use the sub-octave steerable pyramid representation. We tested the sub-octave steerable pyramid with a number of orientations (4, 8, and 16). From this internal analysis, we concluded that the registration algorithm performed best on the sub-octave complex steerable pyramid with 16 orientations filters. We also empirically determined that *α* = 10 generated sufficient amplification to enable accurate registration with minimum artefacts and distortions, and therefore utilized a = 10 for the analyses conducted in this study.

#### 2.4.2 Retrieval of amplified displacements and strain

Extraction of the amplified brain tissue displacement maps from the Damp-MRI and aMRI was achieved using deformable image registration. The domain of deformable image registration algorithms is vast; an excellent overview for medical applications is provided by Sotiras et al.^16^. Given the extensive choices available, we selected the registration algorithm developed by Metz et al.^17^, which was designed for our application (i.e., motion estimation in dynamic medical images). The registration algorithm capitalizes on the following data characteristics/assumptions to extract displacements with sub-voxel accuracy:

1. The data contains a series of images acquired over a short duration in one scan session. Therefore:

a. Monomodal image registration can be performed using a simple dissimilarity metric involving image intensities. The algorithm by Metz et al. minimizes the temporal intensity variance to implicitly align the images to an average reference frame.
b. A group-wise registration approach can be performed, which simultaneously aligns all images in the series. Metz et al.^17^ and Ledesma-Carbayo et al.^18^ both demonstrated that this approach reduces errors in comparison to the pairwise registration approach, wherein each frame of the image series is registered independently to a chosen reference, or consecutively to a neighboring frame. The group-wise approach utilizes information in all images for the registration (leading to a more robust optimization) and also avoids the need to choose a reference frame, which avoids the bias associated with registering to the reference^19,20^.
2. The motion contained in the data series has a physiological source, where the following characteristics are assumed:

a. The physiological displacements were temporally smooth. For the semi-synthetic approach, the reconstructed temporal resolution of the DENSE data is approximately 51 ms, whereas the in-vivo experiment was achieved using a temporal resolution of approximately 13 ms)
b. The displacement fields were spatially smooth. For the semi-synthetic analysis, spatial smoothness was enforced by the smoothing of the DENSE data during the creation of the GTD. For the in-vivo experiment, the analysis was limited to displacements within the brain tissue to reduce the impact of potential discontinuities at the tissue-CSF border.
c. The displacements were temporally cyclic. For both the semi-synthetic and in-vivo experiment, cyclic displacements were enforced by the retrospectively-gated acquisition which was synchronized to the cardiac output.

Extraction of displacements exhibiting spatio-temporal smoothness and cyclic behavior (as described in point 2 above) was achieved in the registration algorithm through the use of a cyclic b-spline transformation model, where the first and last control points on the temporal axis were neighbors^17^.

Registration was performed using Elastix. A multiresolution approach, where the image complexity is reduced by downsampling, or smoothing, can be used to improve registration success. However in this study, we opted to only alter the transformation complexity per resolution, which conserves as much image information as possible for the cost function. This was done by altering the b-spline grid spacing per resolution (in voxels, for [x, y, t]) = [20, 20, 2; 12, 12, 1], where x, y and t denote the 2D and time dimensions. The number of iterations per resolution was set to 1000.

Spatial and temporal sampling was performed to reduce computational complexity. Temporal sampling had the additional benefit of limiting the impact of image artefacts on registration accuracy. Spatial sampling was constrained to voxels within the brain tissue mask. The number of spatial and temporal samples per iteration was set to 2000 and 20, respectively. For the semi-synthetic analysis the use of 20 temporal samples per iteration corresponded to using all available frames in every iteration, whereas for the in-vivo analysis this corresponded to ¼ of all available frames per iteration. The temporal resolution of the in-vivo registration-derived displacements was downsampled to match the GTD resolution.

An average displacement curve for the amplified and ground truth displacements was found by calculating the average displacement within the brain tissue mask for each frame of the cardiac cycle.

2D strain maps were created from the Damp-MRI and aMRI displacement maps, as well as from the GTD, by calculating the 2D divergence of the amplified and ground truth displacements within the brain tissue masks. For each frame, the average strain within the brain tissue mask was calculated, yielding an average tissue strain curve covering the entire cardiac cycle.

#### 2.4.3 Estimation of the amplification factor and measurement agreement

Linear regression was used to estimate the amplification factor (regression slope) and goodness of fit (r^2^) between Damp-MRI derived displacements and the GTD, and also between aMRI-derived displacements and the GTD. For the in-vivo-aMRI, the unsmoothed DENSE measurements were used as the ground truth displacements. To better assess the spatial, temporal and average characteristics of the displacement maps, the linear regression analysis was performed in the following 3 ways:

1. *Between the spatial pattern of the amplified and ground truth displacement maps, for each cardiac frame*. We define the resulting estimated amplification and goodness of fit of this spatial linear regression analysis as EA_s_ and r_s_^2^, respectively. To obtain EA and r_s_^2^, linear regression was first performed on a per frame basis, between voxels in the brain tissue mask of the amplified displacement and GTD maps to determine *f*, the line of best fit. EA_s_ was then defined as the slope of *f*. Secondly, r_s_^2^ was calculated (also on a per frame basis) as shown in Eq. [1], where *ADi* is the Damp-MRI or aMRI-derived displacement in the i^th^ voxel within a brain tissue mask consisting of n voxels. SSr and SSt are the total sum of squares of residuals and the total sum of squares, respectively.

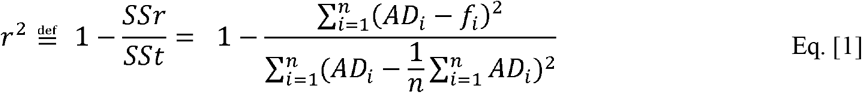

Thus, EA_s_ and r_s_^2^ curves could be constructed, respectively reflecting the estimated amplification and spatial agreement of the amplified displacements to the GTD for each frame of the cardiac cycle.
2. *Between the amplified and ground truth temporal displacement curves*. The estimated amplification and goodness of fit obtained from this temporal regression is defined as EA_t_ and r_t_^2^, respectively, and available for each voxel within the brain tissue mask. EA_t_ and r_t_^2^ maps were constructed through linear regression of the displacement curve of each voxel in the amplified displacement and the GTD maps. In the EA_t_ and r_t_^2^ maps, each voxel represented the estimated amplification and agreement of the amplified displacements at that spatial location over the cardiac cycle, relative to the GTD. For each subject, the mean EA_t_ and r_t_^2^ within the brain tissue mask was calculated.
3. *Between the average displacement curves calculated from the amplified and ground truth displacement maps*. The estimated amplification and goodness of fit obtained from this regression is defined as EA_a_ and r_a_^2^, respectively. To calculate EA_a_ and r_a_^2^, the average AP and FH displacement within the tissue mask was found for each frame of the cardiac cycle, yielding average AP and FH displacement curves for each subject. Linear regression was then performed between the average displacement curves of the Damp-MRI and GTD, and between those of the aMRI and GTD. From this regression, EA_a_ and r_a_^2^ was determined for the AP and FH displacements of each subject.

For all regression analyses, the y-intercept was not forced to zero. The simple percentage error of the Damp-MRI, Dani-aMRI and in-vivo-aMRI amplified displacements relative to the ground truth was also calculated. The percentage error was calculated as 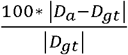, where D and D_gt_ are the amplified and ground truth displacements, respectively.

A similar analysis as above was performed between the amplified strain maps and the ground truth strain to determine the estimated amplification and goodness of fit, thereby evaluating the spatial, temporal and average amplified 2D strain characteristics relative to the ground truth strain.

### 2.5 Statistics

Student’s *t*-tests (paired, 2 tails) were used to assess differences between the estimated amplification and goodness of fit of Damp-MRI and Dani-aMRI derived displacements. Note that a mismatch between Damp-MRI and GTD selectively reflects errors from the registration only, while Dani-aMRI vs. GTD reflects both registration errors and imperfections of the amplification algorithm used by aMRI. Therefore, differences between the estimated amplification of Damp-MRI and Dani-aMRI highlight imperfections arising solely from the aMRI amplification algorithm.

## 3 Results

The analysis of Damp-MRI and Dani-aMRI semi-synthetic data, and the in-vivo-aMRI data were successfully completed on all subjects (n=8). Similar tissue amplification patterns were observed between the semi-synthetic and in-vivo datasets.

Example amplified peak displacement maps and curves for the GTD, as well as for the Damp-MRI and Dani-aMRI derived displacements, are shown in Figure 4. This figure shows that the registration extracted similar displacement fields in all cases, but was most limited for the AP displacements of the in-vivo analysis. The percentage errors of the amplified peak displacements relative to the ground truth are displayed in Figure 5. The group max median percentage errors at peak displacement were (AP, FH, strain): Damp-MRI (44.0%, 44.3%, 81.4%), Dani-aMRI (51.7%, 50.9%, 77.3%), in-vivo-aMRI (145.2 %, 107.2%, 163.9%). Generally, the percentage errors were largest in regions where the ground truth displacements were small.

**Figure 4.**
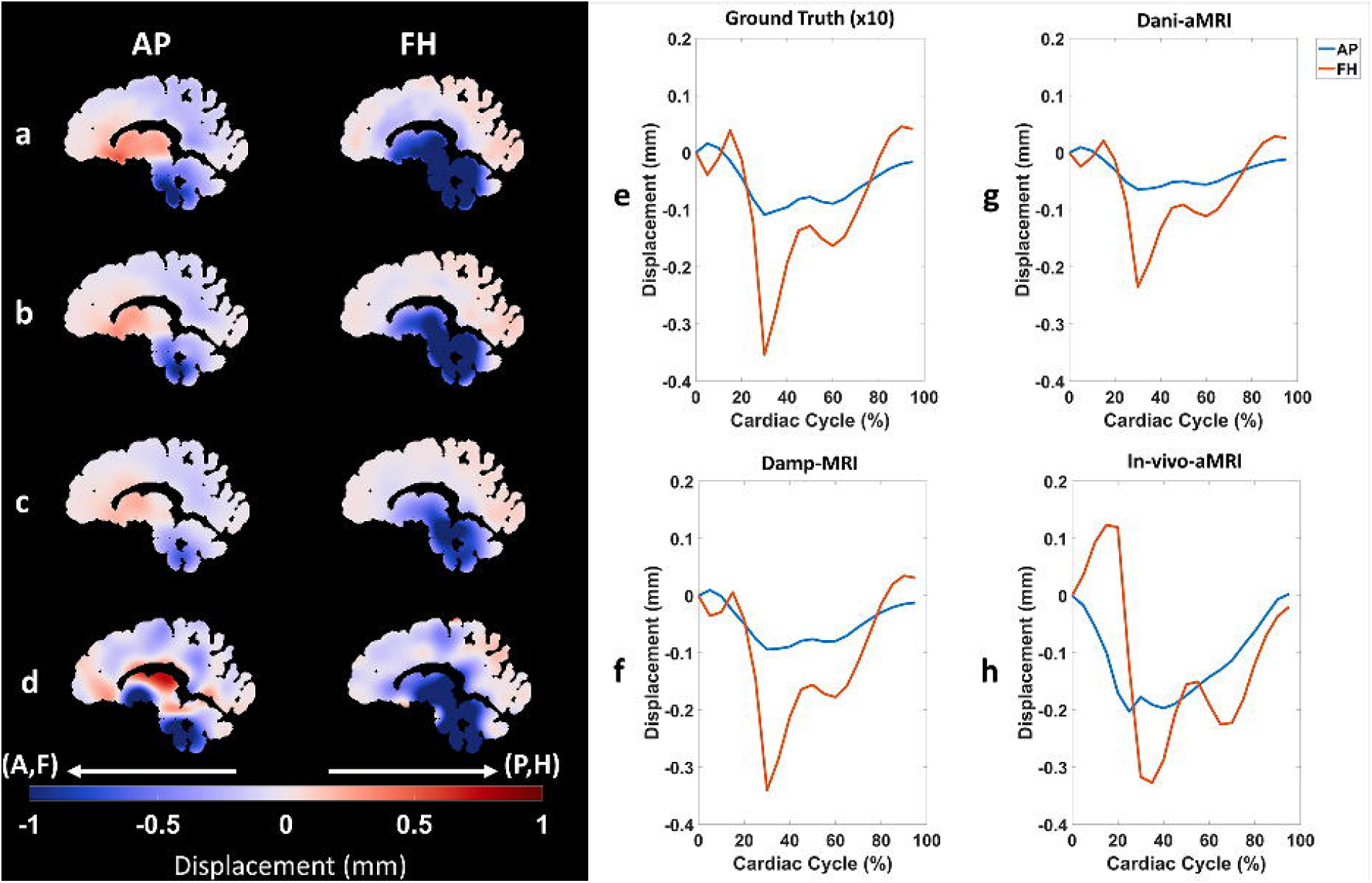
Example AP and FH peak displacement maps derived from Ground Truth (a), Damp-MRI (b), Dani-aMRI (c) and in-vivo-aMRI (d). Ground Truth maps are scaled by 10 to better the fit within the color range of the amplified displacement maps. Maps are shown at the time of peak displacement, relative to triggering at end-diastole. The mean (within the brain tissue) AP and FH displacement over the cardiac cycle for the subject is shown in (e,f,g,h) for Ground Truth (x10), Damp-MRI, Dani-aMRI and in-vivo-aMRI, respectively.

**Figure 5.**
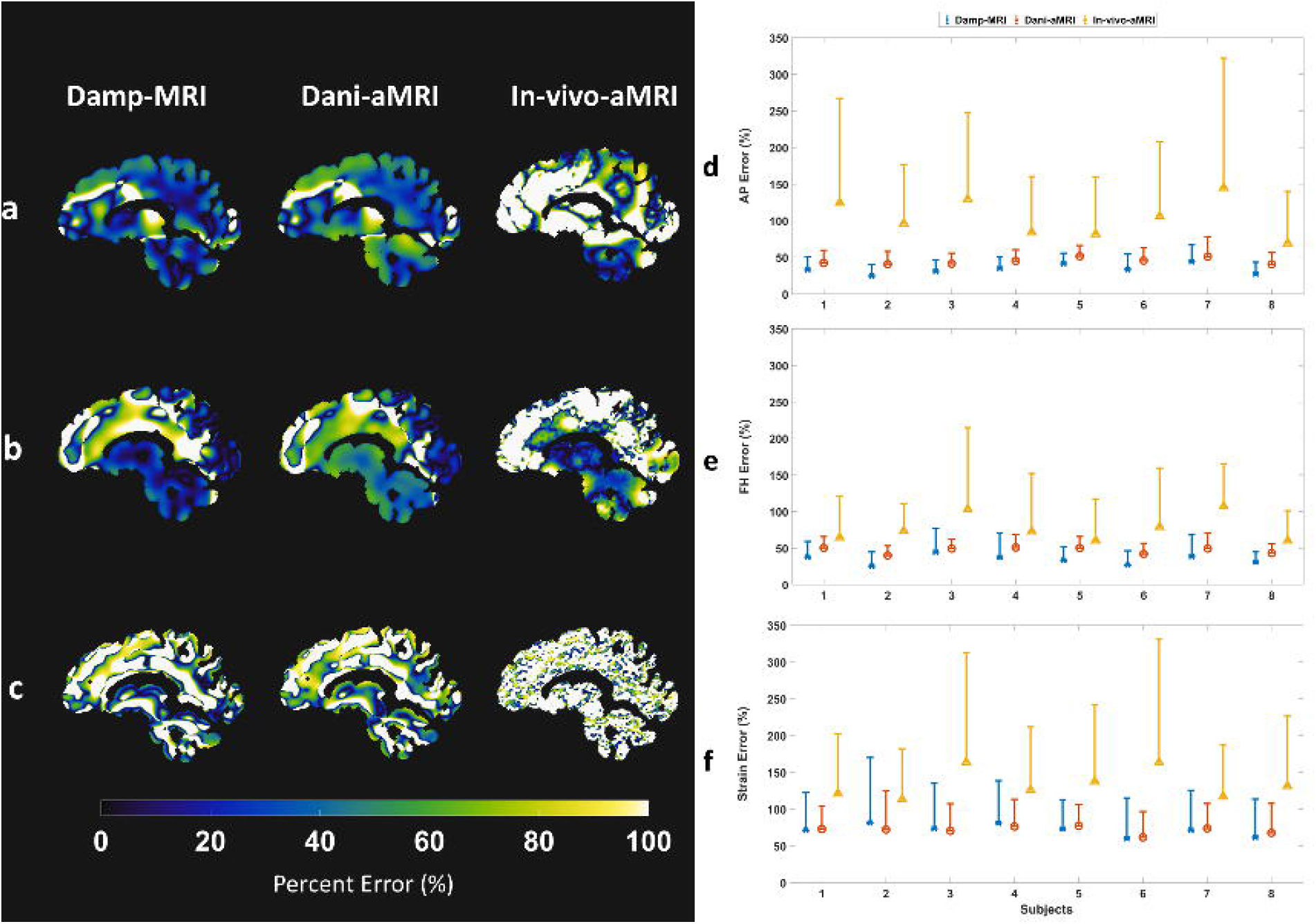
Example AP, FH and strain percentage error maps (a, b, c, respectively) calculated from Damp-MRI, Dani-aMRI and in-vivo-aMRI. Error maps are shown at the moment of peak displacement for the subject. Median percentage error with one-sided interquartile range error bars for AP, FH and strain are plotted for all subjects in d, e, and f, respectively. Percentage error was calculated as 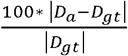, where D and D_gt_ are the amplified and reference displacements, respectively.

Figure 6 shows example EA_t_ and r_t_^2^ maps reflecting the voxel-wise temporal agreement between amplified displacements and the GTD. For the Damp-MRI, large regions of the tissue showed estimated amplification values close to the correct amplification factor of 10. The estimated amplification in these regions was noticeably lower for the Dani-aMRI. The goodness of fit (r_t_^2^) between the Damp-MRI and Dani-aMRI was more similar. Figure 6 (c) and (f) show example EA_t_ and r_t_^2^ maps from the in-vivo-aMRI analysis.

**Figure 6.**
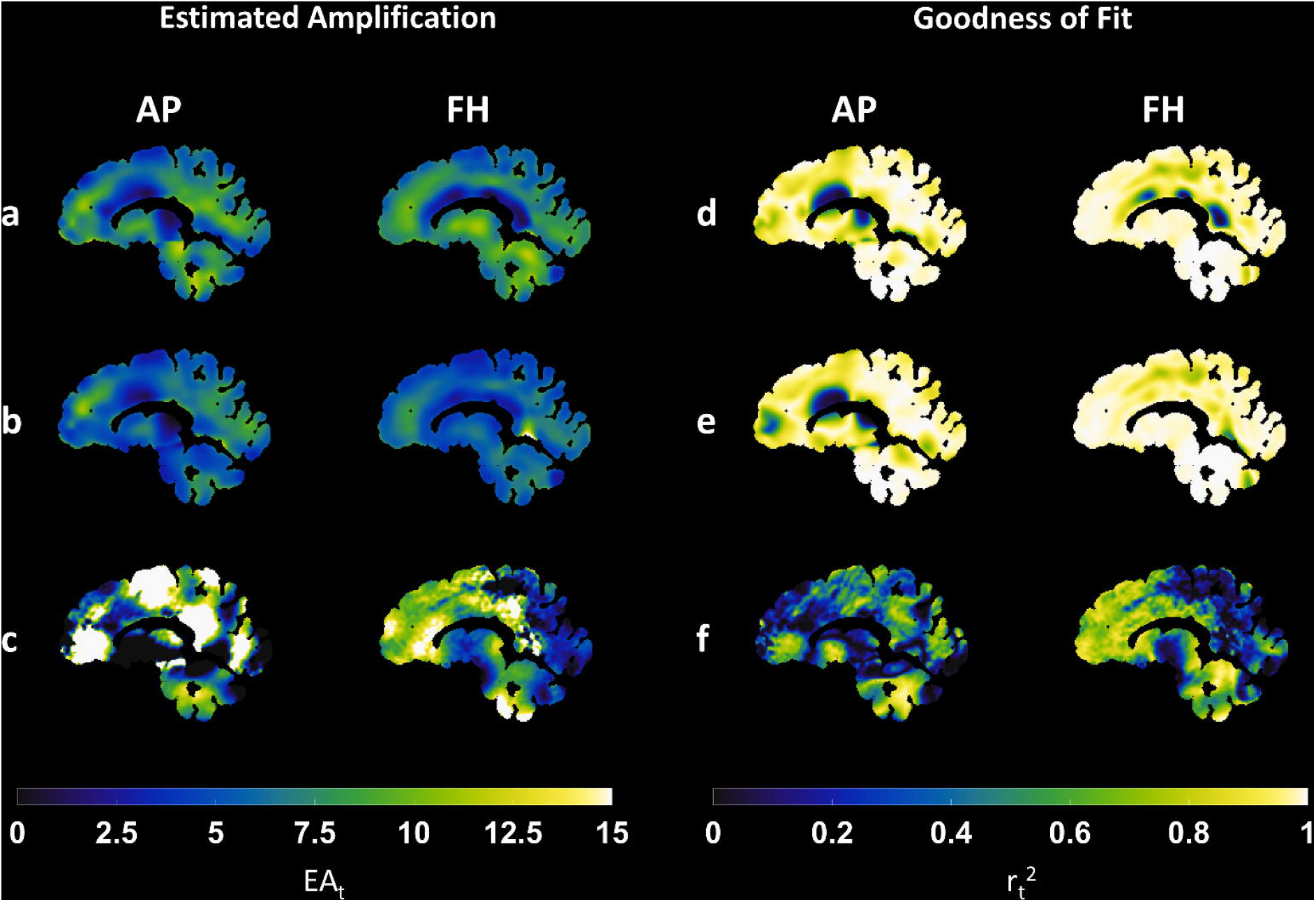
Example Estimated Amplification (EA_t_) and Goodness of Fit (r_t_^2^) maps derived from the temporal analysis of the amplified Damp-MRI (a,d), Dani-aMRI (b,e) and in-vivo-aMRI (c,f) AP and FH displacement maps. In comparison to Dani-aMRI and in-vivo-aMRI, the EA_t_ maps of Damp-MRI appears generally closer to the expected amplification factor of 10, which is desirable.

Figure 7 shows the temporal stability over the cardiac cycle of the EA_s_ and r_s_^2^. The results from the Damp-MRI and Dani-aMRI data were similar, with slightly better EA_s_ and r_s_^2^ results in the former. This shows that errors introduced by aMRI under ideal conditions are limited. The results for both are also noticeably worse at the beginning and end of the cardiac cycle, where the brain tissue displacements are lowest. As with the EA_t_ and r_t_^2^ analysis, accuracy of the extracted displacements from the in-vivo data was limited. This can be concluded from the lower values and higher inter-subject variability in the r_s_^2^ in comparison to that revealed in the semi-synthetic analysis.

**Figure 7.**
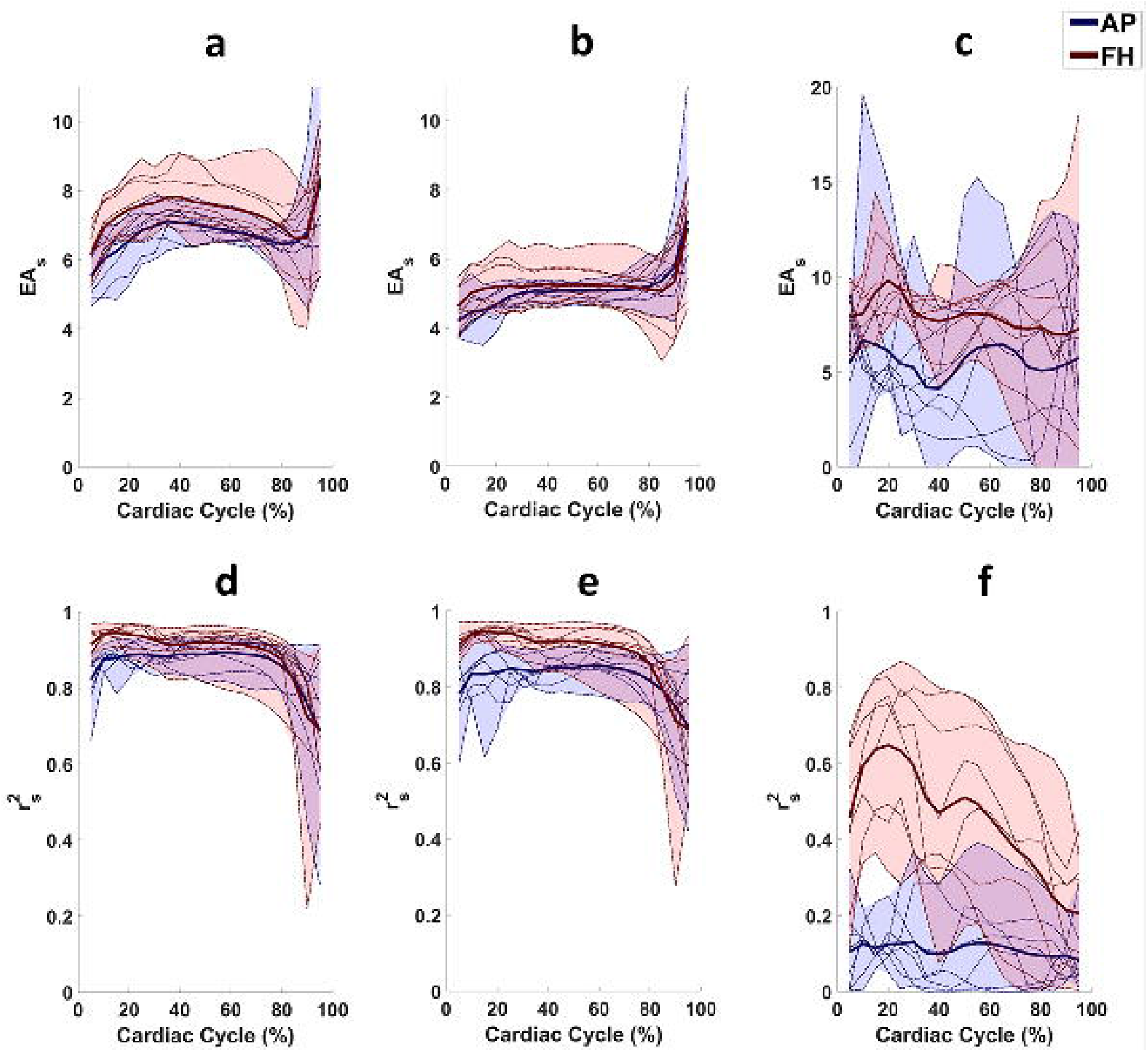
The group mean estimated amplification (EA_s_) and goodness of fit (r_s_^2^) obtained from the spatial analysis for each frame of the cardiac cycle from the Damp-MRI (a,d), Dani-aMRI (b,e), and in-vivo-aMRI (c,f) measurements. The bold red and blue lines represent the group mean value for AP and FH displacements, respectively, whilst the shaded regions reflect inter-subject variability. Displacement offsets resulting from registration were corrected such that the zero displacement coincided with 0% of the cardiac cycle, and is therefore omitted from the plots. The ground truth amplification factor is 10. Note the larger limits used for the y-axis of the in-vivo-aMRI EA_s_ plots.

Strain maps and curves calculated from the displacement maps are shown in Figure 8. The amplitude of the peak strain is much lower for the semi-synthetic analysis relative to the ground truth. However, the general shape of the curves were similar. The general trend showing increased divergence during systole, and subsequent relaxation during diastole was also observed in the strain curves derived from the in-vivo-aMRI data. The percentage error of the strain maps relative to the ground truth are also displayed in Figure 5.

**Figure 8.**
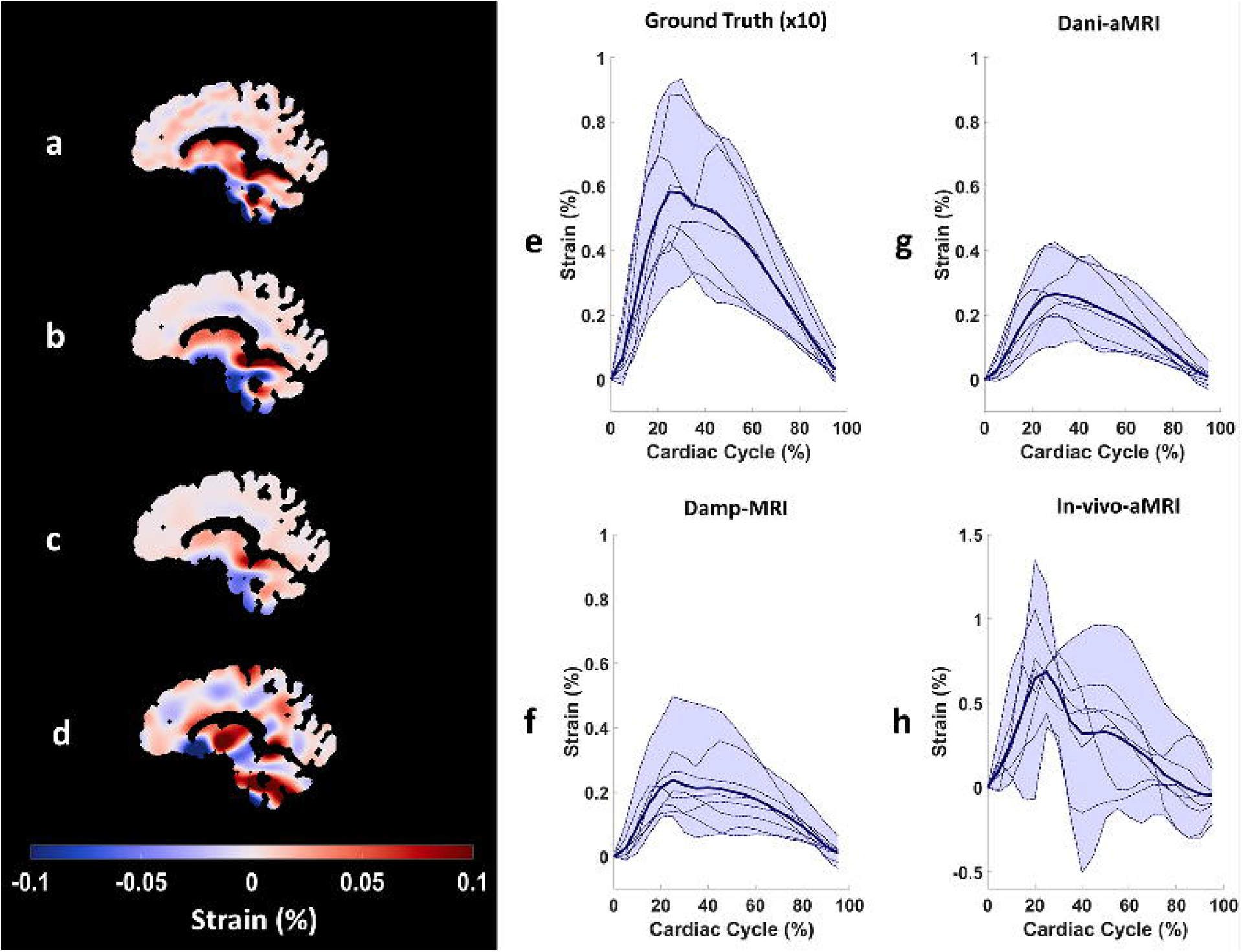
Example peak strain maps derived from Ground Truth (a), Damp-MRI (b), Dani-aMRI (c) and in-vivo-aMRI (d). The Ground Truth map was scaled by 10 to better fit within the color range of the amplified displacement maps. Maps are shown at the time of peak displacement, relative to triggering at end-diastole. The mean (within the brain tissue) strain over the cardiac cycle is shown in (e,f,g,h) for Ground Truth (x10), Damp-MRI, Dani-aMRI and in-vivo-aMRI, respectively. The bold line represents the group mean, and the shaded region the inter-subject variability. Note the larger y-axis limits used for the in-vivo-aMRI strain curves.

Summary results for the group mean estimated amplification and r^2^ are listed in Table 1. The estimated amplification of Dani-aMRI was found to be significantly different than those from the Damp-MRI measurements for both AP and FH displacements, as well as the derived 2-D strain (*P*≤0.05 in all cases). For the spatial regression analysis, only the FH displacements were not significantly different (*P*=0.063). For the temporal regression analysis, only the strain measurements were not significantly different (*P*=0.287). For the average regression analysis, the AP, FH and strain measurements were found to be not significantly different *(P=*0.427, *P*=0.146, *P*=0.085, respectively).

**Table 1.** Group mean estimated amplification (EA) and goodness of fit (r^2^) for spatial, temporal and ROI-averaged analyses for the Damp-MRI, Dani-aMRI and in-vivo-aMRI derived displacements/strain. The true amplification relative to the ground truth displacements/strain was 10. Spatial EA and r^2^ values are reported at peak displacement (end-systole), which for our measurements corresponds to ∼30% of the cardiac cycle.

| | | Estimated Amplification (EA) | | | Goodness of Fit ($r^2$) | | |
| --- | --- | --- | --- | --- | --- | --- | --- |
|  |  | AP | FH | Strain | AP | FH | Strain |
| **Damp-MRI** | Spatial | 6.89±0.62 | 7.55±0.85 | 5.77±1.11 | 0.88±0.03 | 0.92±0.03 | 0.52±0.08 |
|  | Temporal | 6.72±0.46 | 7.34±0.41 | 4.84±0.42 | 0.90±0.02 | 0.96±0.02 | 0.58±0.04 |
|  | Average | 6.89±0.57 | 7.73±0.48 | 3.71±1.22 | 0.97±0.03 | 0.99±0.01 | 0.87±0.12 |
| **Dani-aMRI** | Spatial | 4.92±0.29 | 5.09±0.65 | 3.30±0.75 | 0.84±0.04 | 0.93±0.03 | 0.49±0.07 |
|  | Temporal | 5.53±0.29 | 5.67±0.28 | 3.56±0.33 | 0.88±0.02 | 0.97±0.01 | 0.57±0.04 |
|  | Average | 5.44±0.55 | 5.66±0.36 | 4.59±0.44 | 0.96±0.05 | 0.99±0.00 | 0.97±0.05 |
| **bSSFP-aMRI** | Spatial | 4.44±1.98 | 8.54±1.58 | 0.90±2.06 | 0.11±0.09 | 0.59±0.23 | 0.01±0.02 |
|  | Temporal | 6.95±2.23 | 6.43±2.66 | 0.56±0.84 | 0.39±0.06 | 0.50±0.10 | 0.18±0.03 |
|  | Average | 5.41±5.51 | 7.00±3.25 | 7.21±4.86 | 0.35±0.28 | 0.64±0.28 | 0.56±0.28 |

## 4 Discussion

### 4.1 Summary

We investigated the accuracy of aMRI-derived brain tissue displacements under ideal conditions without image artefacts, using a 2D semi-synthetic dataset with a known amplification factor and known displacements. We found that while the spatial, temporal and average characteristics of brain tissue motion derived from aMRI were very similar to the ground truth, the amplitude was consistently lower than DENSE amplified MRI (Damp-MRI) measurements used for comparison. 2D strain was also computed from the derived displacement fields, where accurate estimation of the amplification factor was found to be more challenging for both Damp-MRI and Dani-aMRI, as reflected by the relatively lower r^2^ values achieved for the strain analyses, in comparison to the displacement analyses. Finally, we investigated the accuracy of aMRI-derived displacements under non-ideal conditions, by comparing the aMRI-derived displacements from in-vivo bSSFP acquisitions to DENSE measurements, which showed relatively poor spatial agreement, particularly for the anterior-posterior displacements.

### 4.2 Agreement

Both Damp-MRI and Dani-aMRI derived displacements showed good agreement to the GTD in the spatial, temporal and ROI-averaged analyses. The performance of Damp-MRI and Dani-aMRI were generally similar, including the inter-subject variation. This shows that deviations from the GTD are mostly related to limitations of the registration algorithm and thus, that the aMRI algorithm performs well under ideal circumstances. These results underscore the importance of the registration algorithm used to extract displacements, and the need for artefact-free images, which could otherwise misguide the registration, as observed in the in-vivo-aMRI experiments. For the strain measurement, r_s_^2^, r_t_^2^ and r_a_^2^ were all lower than those of the AP and FH displacements used in the strain calculation, reflecting the increased uncertainty in the derivation of strain using inaccurate displacements.

### 4.3 Estimated Amplification

The estimation of the amplification factor (regression slope) from Damp-MRI displacements was found to be significantly more accurate than those from Dani-aMRI. Since the Damp-MRI and Dani-aMRI derived displacements were extracted using the same registration algorithm, the difference in estimated amplification can be attributed to the aMRI algorithm. At the same time, recovery of the expected amplification factor of 10 from the Damp-MRI measurements was not achieved, despite the fairly high r^2^ attained in all cases (i.e. r^2^ >0.88). This suggests that intrinsic limitations of the registration approach are at least as important as imperfections of the aMRI algorithm for obtaining accurate extraction of brain tissue displacements. To make aMRI truly quantitative, a calibration process that includes the registration imperfections should be established. Establishing such calibration is not straightforward and might be case-dependent. Nonetheless, even if aMRI is limited to semi-quantitative analyses, there still remains potential for use in cross-sectional or longitudinal clinical research by examining the relative differences in tissue displacements.

Independent of the analysis used, the mean estimated amplification factor was generally larger for FH than AP for both Damp-MRI and aMRI measurements. This may be related to the larger displacements typically observed in the FH direction^10^, and thus tissue displacement in the FH direction may be more clearly identified by the registration algorithm than that displaced along the AP axis. The estimation of the amplification in the 2D strain analysis was found to be less accurate than that of the amplified displacements used in the strain calculation. Intuitively, this can be understood as resulting from the error in the amplified displacements propagating into the strain calculation. Additionally, the different amplification factors found for the AP and FH displacements would further compound the error in the strain calculation, relative to the ground truth.

### 4.4 Linear regression analyses

We utilized linear regression analyses to compare the Damp-MRI and aMRI derived displacements to the GTD, separately exploring their spatial, temporal and average characteristics. Alterations in these characteristics may provide biomarkers for brain tissue disorders in clinical settings^3,12^. Therefore, separate investigation of each characteristic yields a more complete picture of the clinical value of the aMRI-derived displacements, as determined in this study. In particular, poor r_s_^2^ values would indicate that accurate interpretation of the brain tissue strain map on a voxel-wise basis could be challenging since the strain calculation relies on a spatial operation. It should be noted that the percentage errors shown in Figure 5 are a combination of the errors quantified in the spatial amplification and agreement analyses (EA_s_ and r_s_^2^). As an example, regions where the percentage error is 50% could arise simply when the EA_s_ is half of the correct amplification value, even if r_s_^2^ = 1. Similarly, poor r_t_^2^ values would suggest that temporal changes (such as the cardiac-induced swelling of microvasculature) may not be accurately represented in the registration-derived displacements. Therefore, given that aMRI scored highest in the r_a_^2^ analysis, methods involving ROI tissue displacement averaging, such as those utilized by Saindane et al.^3^, may provide useful displacement estimates for aMRI.

### 4.5 In-vivo-aMRI

The displacements extracted from the in-vivo-aMRI images compared poorly to the DENSE measurements, highlighting potential pitfalls of this approach for quantifying brain tissue motion. The mismatch between the DENSE and in-vivo-aMRI derived displacements may have arisen through the presence of bSSFP banding artefacts, which are notably worse at higher field strengths^21^. The registration was only performed using samples derived from within the brain tissue mask, to mitigate the effect of the banding artefacts, which were most noticeable in CSF regions. The extraction of accurate displacements from in-vivo-aMRI with these (amplified) artefacts is nonetheless a challenging registration problem. Both aMRI and the registration algorithm showed excellent performance in the semi-synthetic analysis, even when alternative contrasts and through-plane motion were investigated (see Supporting Information Table S1), suggesting that the image artefacts are the main confounding factor. Similarly to the semi-synthetic approach, poor estimation of the displacements are propagated to the 2D strain calculation, resulting in the low r_s_^2^, r_t_^2^ and r_a_^2^ values observed in this study. Notably the mismatch between the in-vivo derived displacements and the ground truth is a sum effect of registration and aMRI errors, image artefacts and also through-plane motion (although we determined the effect of the latter is limited, see Supporting Information Figure S1).

### 4.6 Clinical Research Value of Brain Tissue Displacement and Strain

The value of brain tissue displacements for clinical research has been demonstrated using integrated PC-MRI velocity measurements^4^, motion tracking software^22^, ultrasound^23^, aMRI^12^, tagged MRI^24^ and DENSE^3^. For the quantification of cardiac-induced brain tissue displacement, it has been argued that DENSE may be an ideal method due to the sensitivity achieved, which is necessary to measure the minuscule amount of tissue motion^5^. Magnetic Resonance Elastography (MRE) is another promising technique to probe brain tissue displacements and (visco)elastic properties^25–27^. These methods typically have a setup complexity rivalling DENSE; whereas DENSE currently requires implementation of a pulse sequence on the MR scanner, MRE can be used in conjunction with the clinically available phase contrast imaging (provided that velocity encoding below 1 cm/s is permitted in the user interface), but nonetheless requires special hardware. However, phase contrast imaging is not ideal for measuring brain tissue motion. This is because peak cardiac induced brain tissue displacements are ∼300 micrometres^5,10^, and the time to peak cardiac output is ∼300 milliseconds^28^, leading to the need for an encoding velocity of ∼1 mm/s. This encoding velocity requires impractically large bipolar gradients, and ultimately leads to the use of a suboptimal encoding velocity, resulting in a large loss in the signal to noise ratio. Additionally, the short gradient ramp times, and large bipolar gradients required to encode the low tissue velocities may lead to eddy currents which yield additional erroneous phase terms to the tissue velocity estimation^29^. Moreover, the stepwise integration of the velocity as a way to yield displacement measures may result in additional errors.

In addition to quantitative measurements of tissue displacement directly available through DENSE, we illustrated that similar to aMRI, DENSE can also provide a qualitative assessment of sub-voxel tissue motion through Damp-MRI, which allows an intuitive visualization of brain tissue physiology. Nevertheless, in this study we sought to establish the potential of aMRI for quantifying brain tissue displacements since it has a relatively high feasibility for widespread clinical availability in comparison to DENSE. The Dani-aMRI derived displacements compared favorably to the Damp-MRI derived displacements, and also to the ground truth. However, the derivation of strain (which may prove to be an important marker for brain diseases^7^) noticeably suffers. We therefore recommend that care be taken regarding the quantification of strain from the amplified displacements using methods as described in this study, as it critically requires knowledge of the amplification factor to correct the resulting amplified strain. Moreover, the results of the in-vivo-aMRI analysis suggests that extreme care must be taken with the MR acquisition protocol used for aMRI analysis of brain tissue motion when applied to real MR images.

### 4.7 Limitations

We explored the performance of aMRI in-vivo using bSSFP cine-images acquired at 7T, where banding artefacts are more severe. Under these non-ideal conditions, the reproducibility of aMRI-derived displacements may also be limited, however this was outside the scope of this study. Additionally, it is conceivable that different slice orientations, positions and thicknesses than that utilized in this study may result in an increased sensitivity to through-plane motion. Therefore, while this work describes the potential and pitfalls of aMRI, future studies should investigate the performance of aMRI algorithms for each particular application, given the illustrated sensitivity to both the image acquisition parameters, as well as the registration methods.

## 5. Conclusion

aMRI-derived displacements are comparable to DENSE under ideal conditions, strengthening the potential of aMRI as a means for investigating brain tissue displacements in the healthy and diseased brain. However, calibration must be established in order to accurately extract displacement (and therefore strain field) maps, otherwise aMRI will be limited to semi-quantitative analyses. The true bottle neck for clinical applications appears to be the artifacts and noise in the in-vivo images, which warrants careful attention on image quality when performing aMRI. Future work is necessary to investigate the extent to which these limitations hinder practical use of aMRI for studying tissue motion in the healthy and diseased brain.

## Supporting information

Supplementary Material for Review and Publication

## List of abbreviations

aMRI: amplified magnetic resonance imaging
bSSFP: balanced steady state free precession
Damp-MRI: DENSE amplified magnetic resonance imaging
Dani-aMRI: DENSE animated amplified magnetic resonance imaging
Dani-MRI: DENSE animated magnetic resonance imaging
DENSE: Displacement Encoding with Stimulated Echoes
EA: Estimated amplification (slope of regression)
GTD: Ground Truth Displacements
T1w: T1-weighted

## Funding

This work was supported by the European Research Council, ERC grant agreement n°337333.

---

Figure S1.Impact of amplified RL motion on sagittal images. a) Anatomical reference normalized to unity. b) Absolute difference of 3D Damp-MRI and 2D Damp-MRI, normalized to unity. The colormap of b) was scaled by a factor of 10 after normalization to better fit within the color range of a), and thus has an unscaled range of [0 0.05]. c) The RL, AP and FH ground truth displacement maps associated with the slice shown in a). Images in b) and c) are shown at the moment of peak displacement.

Table S1.Group mean estimated amplification (EA) and goodness of fit (r^2^) for spatial, temporal and ROI-averaged analyses for the Damp-MRI, Dani-aMRI 2D and 3D derived displacements/strain. The true amplification relative to the ground truth displacements/strain was 10. Spatial EA and r^2^ values are reported at peak displacement (end-systole), which for our measurements corresponds to ∼30% of the cardiac cycle. Please note that the indication of 2D or 3D refers to the motion applied to the synthetic images; the aMRI algorithm is by nature always in 2D, which warrants the analysis of the third, through-plane motion component as performed here.

## Notes

### Competing Interest Statement

The authors have declared no competing interest.

