## Supplementary Material for Review and Publication for "Quantitative comparison between aMRI and DENSE for the assessment of brain tissue motion"

With a semi-synthetic dataset, the performance of aMRI and the registration algorithm under different scenarios was explored. Specifically the impact of through-plane brain tissue motion on the performance of aMRI and the registration algorithm was investigated. The influence of contrast on the registration algorithm was also investigated by utilizing 2D T1-weighted (T1w) images as the reference images, which were deformed by the DENSE, creating T1w Dani-MRI and T1w Damp-MRI (see Figure 1 in the main manuscript). The investigation of through-plane motion provides further insight on the errors observed in the in-vivo analysis.

To investigate the influence of through-plane motion and image contrast on the performance of aMRI and the registration algorithm, semi-synthetic datasets containing T1w contrast were created by first Gaussian smoothing the RL, AP and FH DENSE measurements (kernel = 11 x 11 x 11 voxels, sigma = 1.8), which served as the 3D ground truth displacements (GTD). The smoothed displacements were used to deform the 3D T1w image. 2D slices of the deformed 3D T1w images were then extracted from the same spatial locations as bSSFP acquisitions (section 2.2.2 in the main manuscript), yielding mid-sagittal 2D DENSE animated MRI (Dani-MRI), which exhibit 3D brain tissue motion over the cardiac cycle. DENSE amplified MRI (Damp-MRI) were created by linearly scaling the 3D GTD by a factor of 10 and then using the scaled GTD to deform the 3D T1w. As with the Dani-MRI, 2D mid-sagittal slices were subsequently extracted. As a reference, the same steps were repeated with the RL displacements set to zero, allowing the influence of through-plane motion to be investigated. The analysis steps detailed in section 2.4 of the main manuscript were performed and the results summarized in Supporting Information Table S1 below. Supporting Information Figure S1 contains example images showing influence of amplified RL through-plane motion on the sagittal images.

Minor differences were observed between the 2D and 3D Damp-MRI/Dani-aMRI analyses, suggesting that the impact of through-plane amplified brain tissue motion for mid-sagittal images is limited. The results of the 2D Damp-MRI/Dani-MRI analyses on images with T1w contrast, which were similar to the results obtained from the semi-synthetic bSSFP analyses, supports the hypothesis that these two image contrasts are suitable for use with both aMRI and the registration algorithm.

|  | | **Estimated Amplification (EA)** | | | **Goodness of Fit (r^2^)** | | |
| --- | --- | --- | --- | --- | --- | --- | --- |
|  |  | AP | FH | Strain | AP | FH | Strain |
| 2D Damp-MRI | Spatial | 7.33±0.82 | 8.34±0.70 | 4.17±0.84 | 0.91±0.02 | 0.91±0.03 | 0.29±0.07 |
|  | Temporal | 7.14±0.57 | 8.16±0.48 | 3.74±0.82 | 0.90±0.03 | 0.96±0.03 | 0.46±0.07 |
|  | Average | 7.38±0.44 | 8.46±0.56 | 5.18±1.90 | 0.98±0.01 | 0.99±0.01 | 0.96±0.03 |
| 3D Damp-MRI | Spatial | 6.90±0.94 | 8.12±0.76 | 4.05±0.86 | 0.85±0.08 | 0.90±0.03 | 0.28±0.07 |
|  | Temporal | 7.00±0.91 | 8.25±0.72 | 3.65±0.79 | 0.85±0.04 | 0.95±0.03 | 0.45±0.08 |
|  | Average | 7.45±0.46 | 8.32±0.54 | 6.13±2.23 | 0.97±0.02 | 0.99±0.00 | 0.96±0.03 |
| 2D Dani-aMRI | Spatial | 5.11±0.26 | 5.18±0.60 | 2.41±0.53 | 0.86±0.04 | 0.90±0.03 | 0.26±0.06 |
|  | Temporal | 5.96±0.83 | 5.06±0.90 | 2.39±0.73 | 0.84±0.04 | 0.81±0.04 | 0.45±0.08 |
|  | Average | 5.59±0.53 | 5.02±1.21 | 4.38±1.51 | 0.91±0.03 | 0.86±0.10 | 0.96±0.02 |
| 3D Dani-aMRI | Spatial | 4.99±0.39 | 5.14±0.66 | 2.41±0.57 | 0.82±0.09 | 0.89±0.04 | 0.25±0.05 |
|  | Temporal | 5.90±0.60 | 5.20±1.21 | 2.32±0.71 | 0.80±0.05 | 0.81±0.04 | 0.44±0.08 |
|  | Average | 5.60±0.59 | 4.98±1.22 | 4.54±1.49 | 0.90±0.05 | 0.86±0.10 | 0.95±0.02 |
| **Table S1**. Group mean estimated amplification (EA) and goodness of fit (r^2^) for spatial, temporal and ROI-averaged analyses for the Damp-MRI, Dani-aMRI 2D and 3D derived displacements/strain. The true amplification relative to the ground truth displacements/strain was 10. Spatial EA and r^2^ values are reported at peak displacement (end-systole), which for our measurements corresponds to ~30% of the cardiac cycle. Please note that the indication of 2D or 3D refers to the motion applied to the synthetic images; the aMRI algorithm is by nature always in 2D, which warrants the analysis of the third, through-plane motion component as performed here. | | | | | | | |

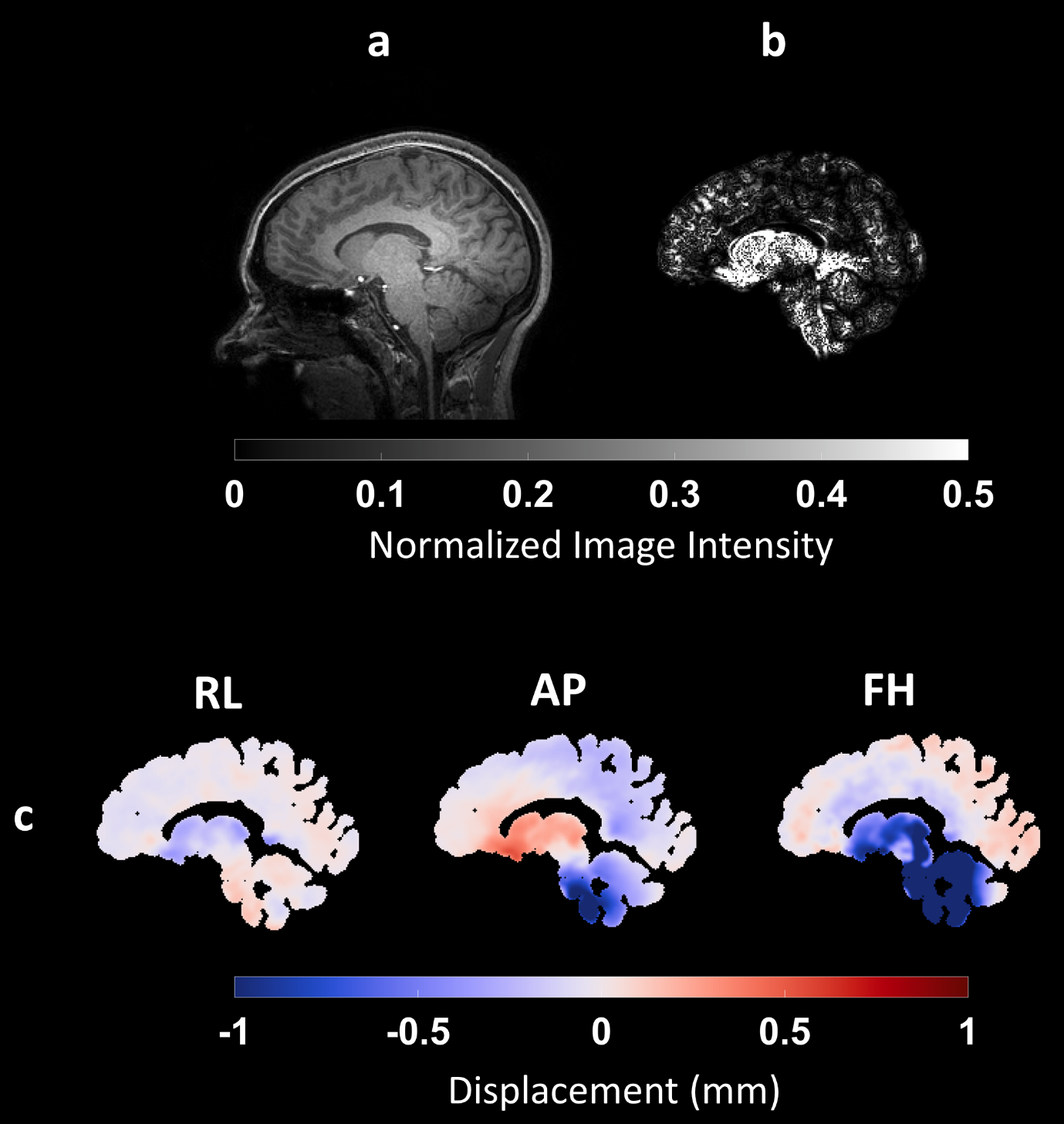

**Figure S1.** Impact of amplified RL motion on sagittal images. a) Anatomical reference normalized to unity. b) Absolute difference of 3D Damp-MRI and 2D Damp-MRI, normalized to unity. The colormap of b) was scaled by a factor of 10 after normalization to better fit within the color range of a), and thus has an unscaled range of [0 0.05]. c) The RL, AP and FH ground truth displacement maps associated with the slice shown in a). Images in b) and c) are shown at the moment of peak displacement.
